# Modeling the Co-Activation of Inflammatory Genes Mediated by NFκB and IRF-3 During Viral Infections

**DOI:** 10.1101/2025.05.13.653653

**Authors:** Syona Tiwari, Soumen Basak, Rakesh Pandey

## Abstract

Innate immune responses of host cells are activated upon exposure to external stimuli such as viral infections. These activations trigger the signaling pathways involved in antiviral responses. Activated pathways play a crucial role in the regulation of inflammatory and antiviral responses involving transcription factors, cytokines, cell surface viral sensors, and the activation of target genes. Although the role of key transcription factors such as NF-*κ*B and IRF-3 in the regulation of antiviral genes that encode type I interferons (IFN*α/β*) is identified and well explored. The precise mechanistic understanding that governs their complex context-dependent temporal activation remains elusive. Here, we have developed a mathematical model to demonstrate the co-regulation of the expression of the IFN*β* gene by Nuclear Factor kappa B (NF*κ*B) and IFN Regulatory Factor-3 (IRF-3) after their activation in host cells by the viral analogue Poly I:C. In our model, the binding activities of the transcription factors NF*κ*B and IRF-3 at the IFN*β* promoter are modeled using a probability-based framework. Our results demonstrate that the expression pattern of IFN*β* postviral infections would be pertaining to the co-activation of that by NF*κ*B and IRF-3. In addition, the immediate expression of the IFN*β* gene and, thus, the initial host response would be mainly attributed to immediate activation of NF*κ*B while IRF-3 would be responsible for delayed but persistent expression of the IFN*β*. Our model provides an underlying molecular mechanism for a similar response observed for Newcastle disease virus and the Vesicular stomatitis virus infection. Furthermore, we suggest that the rapid as well as an elevated and persistent expression of IFN*β*, would be observed later due to a temporal bias in binding of both transcription factors NF*κ*B and IRF-3, highlighting the need for coregulation of IFN*β* by two different signaling pathways in a sequential manner. Also, our model predicts that the binding affinity of IRF3 to the promoter region of IFN*β* would be equal to or larger than that of NF*κ*B for a strong and persistent antiviral responses.

## 1 Introduction

The innate immune response is the first line of defense of hosts cells against invading pathogens. It is initiated after the identification of viral or bacterial nucleic acids through a pathogen-specific host receptor. This initial immune response activates a cascade of cell signaling pathways that target the containment and elimination of the pathogen. An important aspect of this response involves the activation of pro-inflammatory and antiviral transcription factors, such as NF*κ*B, AP-1, NF-IL6, NFAT, IRFs, and STATs, which regulate the expression of Interferons(IFNs) [1, 2].

IFNs are cytokines that suppress viral replication in host cells by transcriptional induction of various IFN-stimulated genes (ISGs) [3]. Typically, three types of IFNs (Type I, II, and III) are believed to play a role in activating the antiviral state of infected cells. [4]. Of these, Type I interferons (IFN *α*/*β*) and their receptors are expressed by most cells that initiate the immune response in the early stage and immunosuppression during chronic infection [5]. Type II IFN expression is mediated by a single gene (IFN *γ*) in T cells or natural killer cells (NK) that occurs in all cells expressing its receptors. However, Type III IFNs(IFN *λ*1, IFN *λ*2, and IFN *λ*3) are produced by specific cell types and therefore cause an acute response to viral infection in cells (such as the formation of epithelial barriers) [6]. Respiratory viral infections, such as SARS-CoV-2, pose a high risk to these barrier cells present in the respiratory tract, thus activating the targeted action of Type III IFN [7].

The induction and expression of IFN and its associated genes are the result of several molecular mechanisms and signaling pathways. Of which, the inflammatory signaling pathway mediated by the transcription factor NF*κ*B and the receptor-mediated retinoic acid-inducible gene I (RIG) signaling pathway that involves the IRF-3 transcription factor are significant in the antiviral mechanism. NF*κ*B is observed in its nascent form, that is, the inhibitor of kappa B (I*κ*B) is attached to it in the absence of any external stimulus, such as nucleic material from the invading pathogen. However, an external stimulus to the host cell by viruses (such as the viral analog Poly I:C) or Lipopolysaccharide (bacterial analog) leads to the activation of the NF*κ*B pathway.

After the entry of the viral genome into the cytoplasm, they are sensed by pattern recognition receptors (such as Toll-like receptor (TLR), RIG-like receptor(RLR), etc.). This leads to phosphorylation of I*κ*B*α* bound to NF*κ*B, which is then degraded in the cytosol. I*κ*B*α* is therefore disassociated from the NF*κ*B subunit, thus activating it [8]. Active NF*κ*B transcription factor is then translocated within the nucleus. There, it binds to the promoter site of inflammatory genes such as Type I IFNs (IFN *α*/*β*) and regulates their expression [9, 3].

The other signaling pathway that is activated upon recognition of non-self RNA (such as viral RNA) by the RIG-I receptor encodes the transcription of interferon regulatory factors (IRFs). Ingression of double-stranded RNA (dsRNA) in the cytosol leads to the specific activation of one of the IRFs, i.e., the IRF-3 signaling pathway [10, 11]. The entry of RNA virus into the cytoplasm signals the adapter protein Mitochondrial Antiviral Signaling Protein (MAVS) to form a complex with RIG and propagate the signal to downstream molecules. Poly I:C is a synthetic analog of dsRNA that is used to mimic viral infections that produce dsRNA in their replication cycle. Since Poly I:C is mainly associated with RNA viruses, it has been suggested that it may also be relevant to study the immune response to SARS-CoV-2 infection [12]. For this purpose, an infection by the viral analog Poly I:C initiates signaling through pattern recognition receptors (PRR) such as RIG I [13]. The signal propagated by the Poly I:C-RIG I-MAVS axis recruits the TNF-Receptor-Associated Factor (TRAF) protein towards itself [14]. This TRAF protein relays the activation of the IRF-3 and NF*κ*B signaling pathways through TANK-binding kinase (TBK) [15]. TBK is phosphorylated into its active form that subsequently phosphorylates specific serine residues on IRF-3 [16]. After the phosphorylation of IRF-3, it dimerizes and then localizes inside the nucleus. Here, it functions as a crucial transcription factor for binding to the promoters of type I IFNs (IFN *α*/*β*), several cytokines, as well as several antiviral ISGs [17].

Simultaneous activation of both IRF-3 and NF*κ*B signaling pathways transcribes IFNs (IFN *β*) that encode the antiviral state in the host cell. The activation of such a state is believed to lead to rapid degradation of viral nuclei, inflammation in the host, and activation of IFN-induced genes and gene products [18, 19, 20, 21, 22, 23, 24, 25, 26].

Expression of transcription factors NF*κ*B and IRF-3 to pathogenic stimulus and their biochemical mechanisms have been modeled previously [27, 28, 29, 30, 31, 32, 33]. A mathematical model using the stochastic framework has previously been studied to model the regulation of IFN *β* by these transcription factors [28]. In this model, three inputs to the system (Poly I:C, IFN *β*, and LPS) trigger the immune signaling pathways and their downstream effects on gene expression. Here, three transcription factors, STAT 1/2, NF*κ*B, and IRF-3, are modeled as regulatory systems that are activated after stimulation with Poly I:C or LPS.

However, a model for the co-regulation of Type I IFNs, specifically by NF*κ*B and IRF-3, is still elusive. Also, a model to calculate the promoter binding activity of IFNs coregulated by NF*κ*B and IRF-3 is less explored.

In the present study, we model the co-regulation of IFN *β* in response to the Poly I:C input (viral analog) mediated by NF*κ*B and IRF-3 in the host’s cell through an ODE-based deterministic model. To model both signaling pathways, we incorporated the established mechanisms of the activation of NF*κ*B signaling pathway into our model [27]. The IRF-3 signaling pathway was modeled based on known biochemical interactions [28]. We calculated the expression of IFN*β* after binding both the transcription factors to the promoter site of the IFN *β* gene using a Hill-like equation. We varied the parameters of the IRF-3 module to two and three orders of their magnitude and studied its effect on the expression of IRF-3 and IFN *β* inside the nucleus of the host’s cell.

Our model results demonstrate that for early and strong as well as a delayed but persistent antiviral and inflammatory response, binding of both NF*κ*B and IRF-3 is required. In addition, the early immune response to viral infection would be attributed to the activity of transcription factor NF*κ*B, whereas the transcription factor IRF-3 would be responsible for sustained albeit delayed IFN*β* expression. This result is consistent with an observation for Newcastle Disease Virus (NDV) and Vesicular Stomatitis Virus (VSV) [34]. We propose that it occurs due to a temporal bias in the binding of NF*κ*B and IRF-3. However, binding of both transcription factors is required. This hypothesis is in line with the finding of Cheng et al, [35] and proposed the function of both transcription factors as a “Sequential OR gate”. Furthermore, our model predicts that the binding affinity of IRF3 would be higher than that of NF*κ*B for an antiviral response observed experimentally.

## 2 Model Development

We simulate the host’s innate immune response to viral infection by modeling the biochemical reactions involved in the activation of NF*κ*B and IRF-3 signaling cascades using an ODE-based model (Figure 1). Poly I:C, a viral analog, acts as an input to the model, resulting in the activation of NF*κ*B and IRF-3 signaling pathways upstream of IFN*β*. Here, the dynamic expression profile of IFN*β* is computed through the numerical solution of the ODEs for the nuclear concentration of the transcription factors, followed by the calculation of promoter binding activity of respective transcription factors to the host’s DNA.

**Figure 1:**
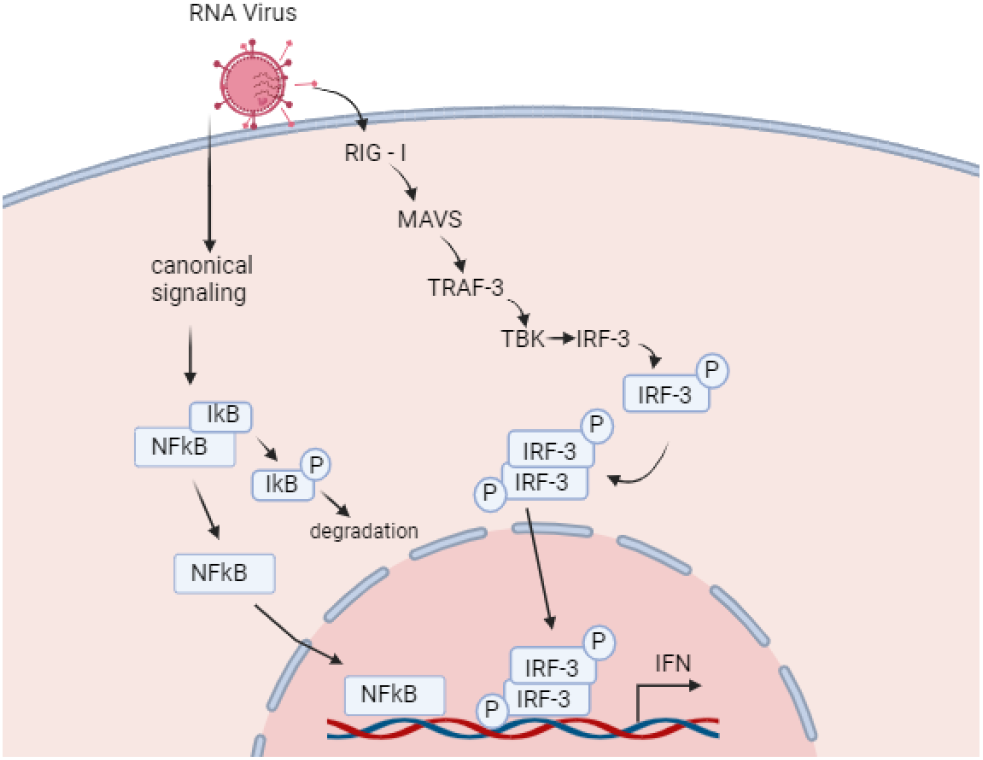
A schematic representation of the NFκB and IRF-3 signaling pathways leading to the activation of Type I IFNs in response to viral infection. Here, activation of both the signaling pathways is assumed through the same input Poly I:C

### Modeling of the NF*κ*B Pathway

NF*κ*B has been comprehensively explored for its role in the inflammatory response across different cells [36, 37, 38, 27, 39, 40]. The canonical and non-canonical pathways of NF*κ*B’s activation have been mathematically modeled earlier [27]. In our model, the canonical pathway is given a Poly I:C input upstream of the level of NEMO activation, and its time-dependent value is determined by its half-life and initial concentration. The non-canonical or the alternative signaling of the NF*κ*B pathway is attenuated by giving a null value to the NIK input. The nuclear concentration of active NF*κ*B, in response to a viral stimulus, is subsequently plotted using MATLAB Software [41].

### Modeling of IFN Regulatory Factor - 3 Pathway

Previous studies have introduced several mathematical models that utilize stochastic [28], and deterministic [42, 43, 44, 45, 46, 47, 48] mathematical modeling framework to model the IRF-3 signaling pathway as a part of their methodology. We model the IRF-3 signaling pathway after surveying the literature for all the biochemical species involved in the sensing of viral nuclear material, transduction of its signal, and activation of IRF-3 transcription factor [28] (Supplementary Information: Table 1 and Table 2). To model the activation of IFN *β* downstream of the IRF-3 pathway upon binding of IRF-3 transcription factor to the promoter region, we provide Poly I:C input to the RIG-mediated activation of IRF-3 arm as illustrated in Figure 1. MATLAB Software is used to compute the numerical solution of the ODE systems involving all the biochemical species modeled of the IRF-3 pathway (Figure 2). The same input signal of POLY I:C (viral analog) is given to the IRF3-arm that was given to the NF*κ*B arm, and the levels of phosphorylated nuclear dimer of IRF-3 are calculated. Values of 24 kinetic parameters (rates reported in *min*^*−*1^) are assumed in the model and the physiological range of them were obtained from literature [48, 28, 49]. Values of these parameters are guessed by varying their values to the two and three orders of their magnitude (Figure 3, Supplementary Information: Figure 1).

**Table 1:**
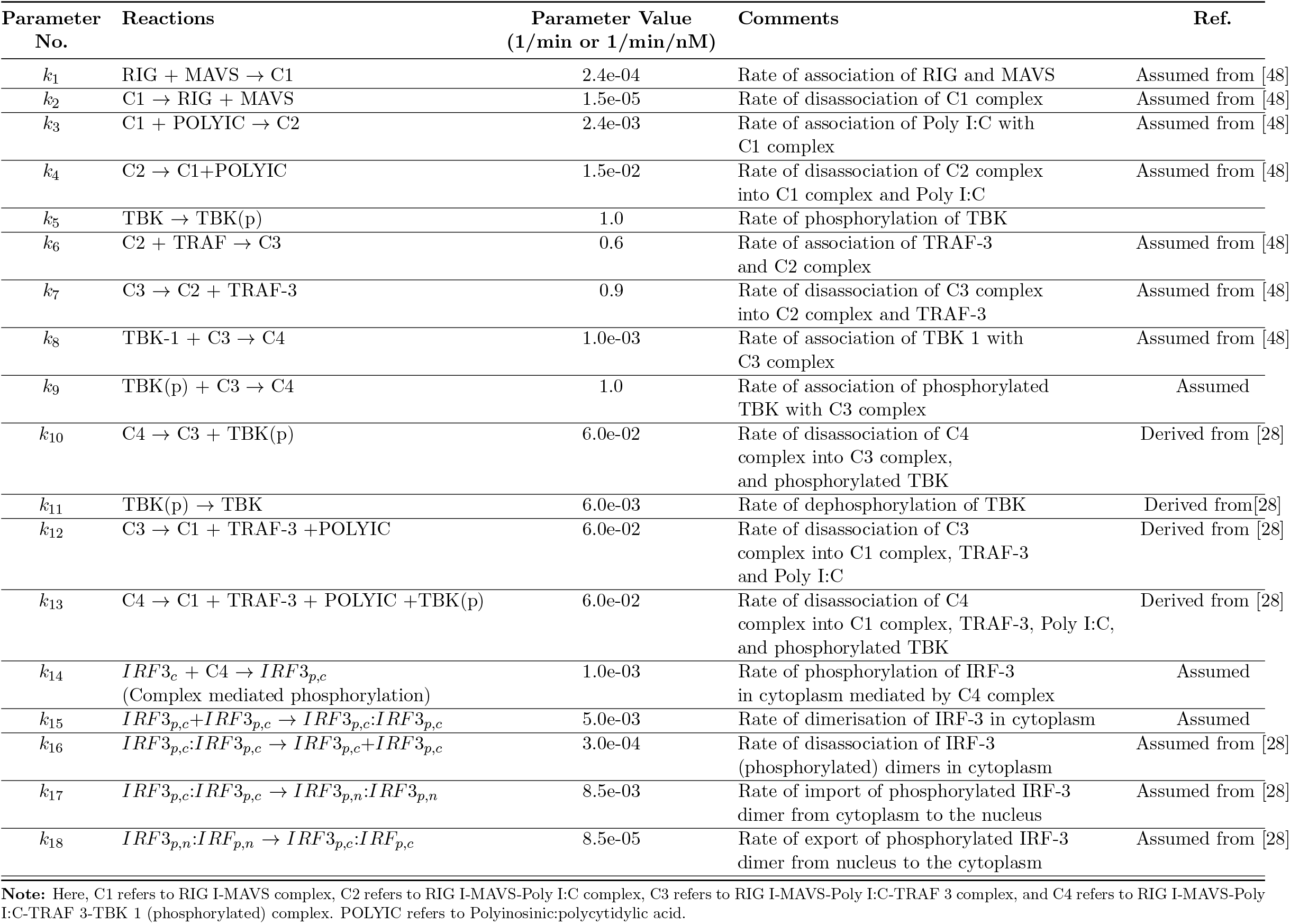
List of biochemical reactions in the IRF-3 Model and description of the model parameters.

**Table 2:** List of biochemical reactions in the IRF-3 Model and description of the model parameters.

| Parameter No. | Reactions | Parameter Value<br>(1/min or 1/min/nM) | Comments | Ref. |
| --- | --- | --- | --- | --- |
| $k_{19}$ | $IRF3_{p,c} \rightarrow IRF3_{p,n}$ | 8.5e-03 | Rate of import of phosphorylated IRF-3 monomer from cytoplasm to the nucleus | Assumed from [28] |
| $k_{20}$ | $IRF3_{p,n} \rightarrow IRF3_{p,c}$ | 8.5e-05 | Rate of export of phosphorylated IRF-3 monomer from nucleus to the cytoplasm | Assumed from [28] |
| $k_{21}$ | $IRF3_{p,n}:IRF3_{p,n} \rightarrow IRF3_{p,n} + IRF3_{p,n}$ | 3.0e-04 | Rate of disassociation of phosphorylated IRF-3 dimer into its monomers (inside the nucleus) | Assumed |
| $k_{22}$ | $IRF3_{p,n} + IRF3_{p,n} \rightarrow IRF3_{p,n}:IRF3_{p,n}$ | 5.0e-04 | Rate of association of phosphorylated IRF-3 into its dimers inside the nucleus | Assumed |
| $k_{23}$ | $RIG \rightarrow \phi$ | 2.1e-07 | Rate of degradation of RIG | Assumed from [49] |
| $k_{24}$ | $MAVS \rightarrow \phi$ | 2.2e-07 | Rate of degradation of MAVS | Assumed from [49] |
| $k_{25}$ | $TBK-1 \rightarrow \phi$ | 1.2e-07 | Rate of degradation of TBK-1 | Assumed from [49] |
| $k_{26}$ | $TRAF-3 \rightarrow \phi$ | 6.9e-09 | Rate of degradation of TRAF-3 | Assumed from [49] |
| $k_{27}$ | $IRF3_c \rightarrow \phi$ | 8.8e-08 | Rate of degradation of IRF-3 (in cytoplasm) | Assumed from [49] |
| $k_{28}$ | $IRF3_{p,n} \rightarrow \phi$ | 8.8e-08 | Rate of degradation of phosphorylated IRF-3 monomer (in nucleus) | Assumed from [49] |
| $k_{29}$ | $IRF3_{p,c}:IRF3_{p,c} \rightarrow \phi$ | 8.8e-09 | Rate of degradation of phosphorylated IRF-3 dimer (in cytoplasm) | Assumed from [49] |
| $k_{30}$ | $IRF3_{p,n}:IRF3_{p,n} \rightarrow \phi$ | 8.8e-09 | Rate of degradation of phosphorylated IRF-3 dimer (in nucleus) | Assumed from [49] |
| $k_{31}$ | $\rightarrow RIG$ | 0.5 | Basal rate of formation of RIG | Assumed |
| $k_{32}$ | $\rightarrow MAVS$ | 0.5 | Basal rate of formation of MAVS | Assumed |
| $k_{33}$ | $\rightarrow TBK-1$ | 0.5 | Basal rate of formation of TBK-1 | Assumed |
| $k_{34}$ | $\rightarrow TRAF-3$ | 0.5 | Basal rate of formation of TRAF-3 | Assumed |
| $k_{35}$ | $\rightarrow IRF3_c$ | 0.5 | Basal rate of formation of IRF-3 in cytoplasm | Assumed |

**Figure 2:**
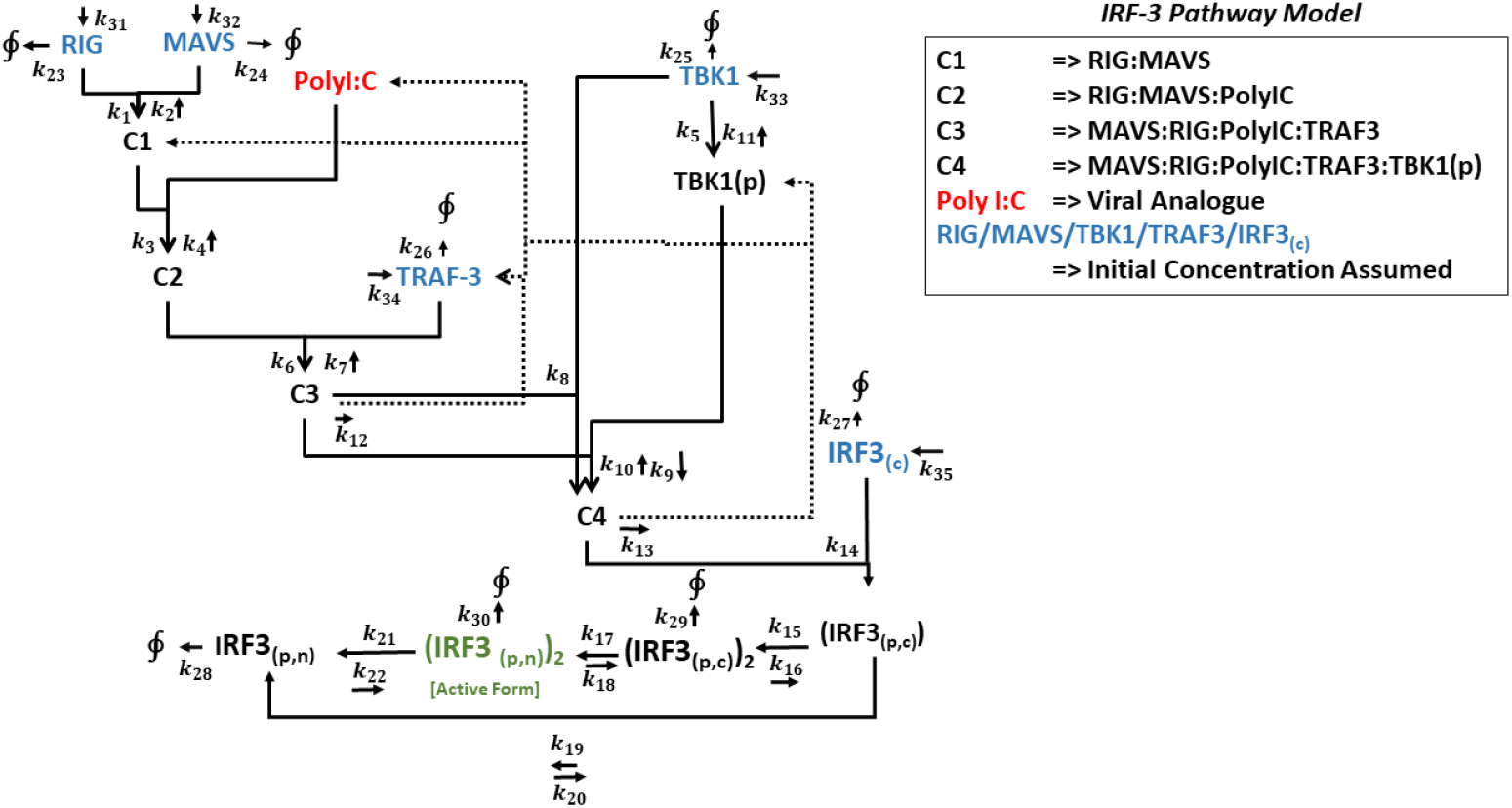
A circuit of all the biochemical interactions in the IRF-3 signaling pathway that has been modeled in our system of equations. Here, the complexes formed during the reactions are represented by C1, C2, C3, and C4. Default values of the parameters have been summarized in Table 1 and Table 2. Poly I:C (in red) is a viral analogue that triggers the activation of the IRF-3 signaling cascade. An initial concentration for the species (in blue) RIG, MAVS, TRAF-3, TBK 1, and inactive IRF-3 in the cytoplasm has been assumed. The IRF-3 species (in green) represents the active nuclear dimer of phosphorylated IRF-3 that functions as a transcription factor.

**Figure 3:**
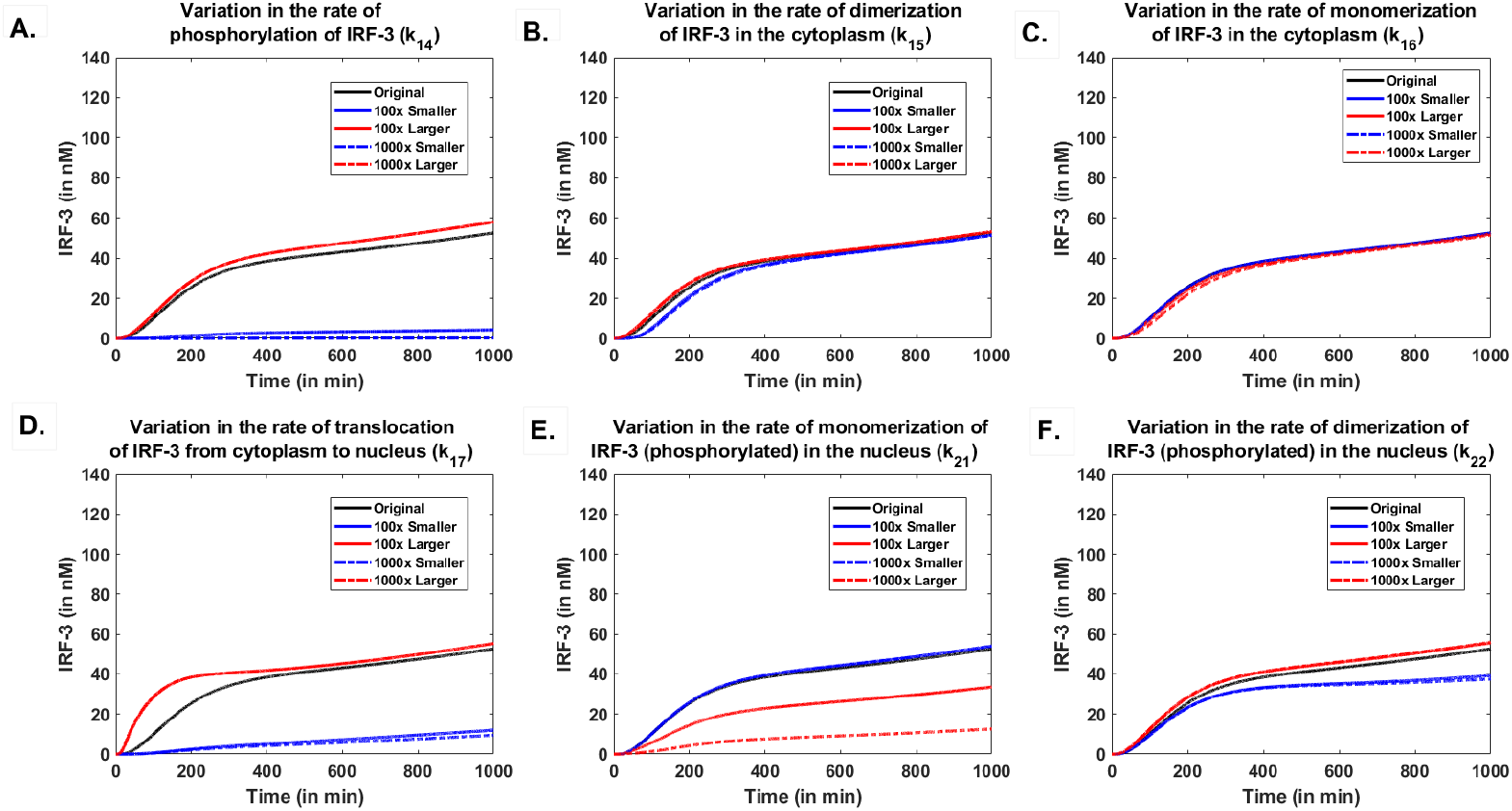
Systematic Parameter Analysis: Effect of varying model parameters (k_14_ to k_19_) on the dynamics of nuclear concentration of active IRF-3 over time has been shown. Here, the values of the respective parameters were increased or decreased by 2- or 3-fold (represented by red and blue curves) of their original values (represented by black curves). Poly I:C Input I, with a half-life of 20 min and initial concentration of 500 nM, was used as input for the present parameter analysis in the model.

### Modeling of the Coregulation of IFNs

Type I IFNs are activated by both NF*κ*B and IRF-3 signaling pathways. They co-regulate the IFNs inside the nucleus upon binding of the respective transcription factors (i.e. NF*κ*B and IRF-3) on the promoter binding site of the host cell’s DNA. We use the following mathematical modeling framework using Hill-like equations to model the effect of activators on the transcriptional response of the gene of interest that is proposed elsewhere [50, 51, Appendix A].

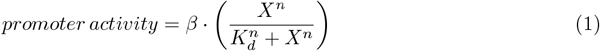

Here, ‘*β*’ represents the maximal transcription rate, ‘X’ represents the concentration of transcription factor binding to the promoter site, ‘*K*_*d*_’ represents the disassociation constant, and ‘n’ represents Hill coefficient like parameter for the co-operativity of binding.

The binding of NF*κ*B and IRF-3 to the respective binding sites upstream to the IFN*β* are modeled through the following equation. Since both the pathways are induced by the viral analog, there would be the presence of both the transcription factors inside the nucleus, and therefore the transcriptional response of IFN*β* would be modeled as follows.

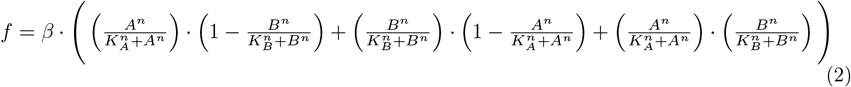

here ‘A’ and ‘B’ represent the nuclear concentrations of the transcription factors IRF-3 and NF*κ*B, respectively. ‘*K*_*A*_’ and ‘*K*_*B*_’ represent their dissociation constants (defined as the concentration of the respective transcription factor for half-maximal binding to the promoter site) and *n* is Hill coefficient that represents the co-operativity of transcription factors to the DNA binding site as described earlier in [52]. In equation 2, three terms are considered to quantify the transcribed mRNA level of the IFN*β* gene. The first term 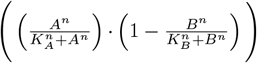 quantifies the probability of binding of the transcription factors to the promoter region when NF*κ*B is bound and IRF-3 is not, the second term 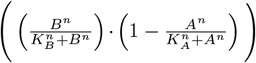 represents the probability when IRF-3 is bound and NF*κ*B is not, and the third term 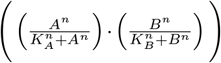 represents the probability of a scenario when both NF*κ*B and IRF-3 are bound to their respective binding sites.

### Viral Analog Input Signal: Poly I: C

Poly I:C, the viral analog of double-stranded RNA, is a synthetic mimic of viruses that is administered in vitro through transfection in the host’s system. Previous studies have determined its half-life in the human serum through radiochemical analyses[53]. In our model, we provided two Poly I:C signals using an initial concentration of 500nM and 50nM, with a half-life of 20 *min*^*−*1^ and 6 *min*^*−*1^, respectively as represented in Figure 4A.

**Figure 4:**
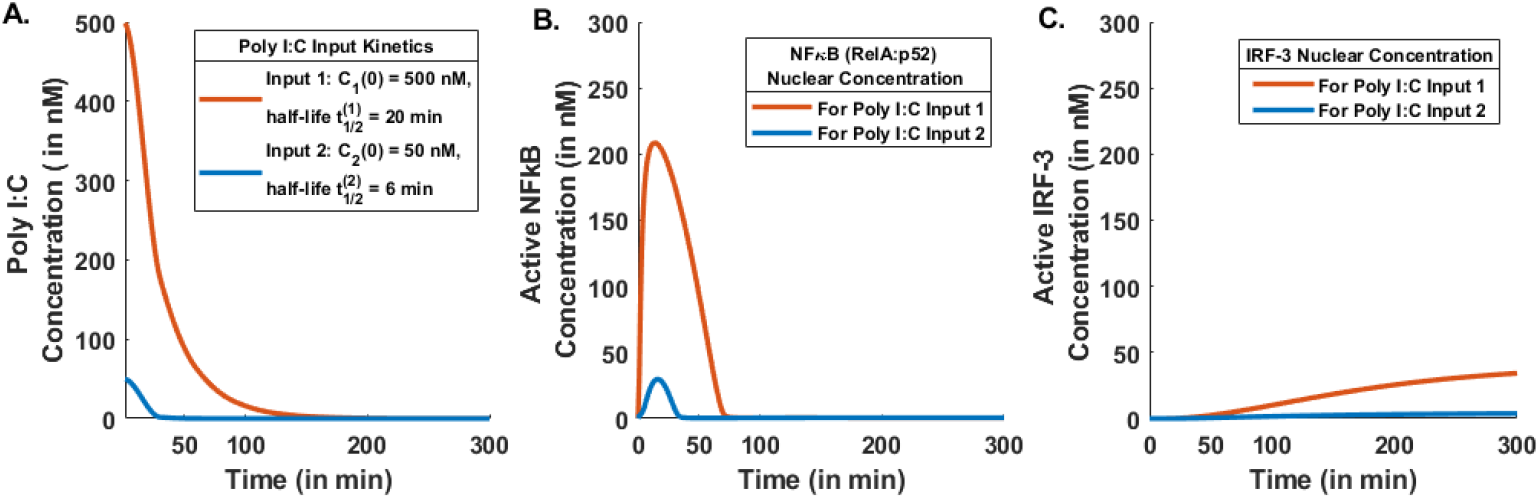
Dynamic response of the transcription factors NFκB and IRF-3 to a common input signal Poly I:C (A.)Poly I:C Input Kinetics. The viral RNA analog Poly I:C is administered through two independent inputs modeled as exponential decay. Here, Input 1 (orange curve) represents the initial concentration C_1_(0) = 500 nM and half-life 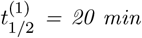, and Input 2 (blue curve) represents the initial concentration C_2_(0) = 50 nM and half-life 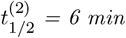.**(B.)** Active and nuclear expression levels of NFκB (in nM) in response to Poly I:C Input 1 (orange curve) and Poly I:C Input 2 (blue curve).**(C.)** Active and nuclear expression levels of IRF-3 in response to Poly I:C Input 1 (orange curve) and Poly I:C Input 2 (blue curve).

## 3 Results

### 3.1 Response of IRF-3 Arm and Role of Model Parameters

For the NF*κ*B arm of signaling, we follow an ODE model developed elsewhere ([27]) while for IRF-3 we have developed an ODE-based deterministic model. Using our model for IRF-3 we can calculate the levels of active IRF-3, i.e. phosphorylated dimer of IRF-3 inside the nucleus.

Since the values of the model parameters (described in Tables 1 and 2) are unknown, we first assume a default value for them and later vary them by 2 to 3 orders of magnitude to explore the effect of systematically varying the model parameters on the level of active IRF-3. For this exploration, we assume that the IRF-3 arm is stimulated through an input signal (Poly I:C) with a half-life 20 minutes. Our results suggest that the concentration of nuclear active IRF-3 would not change significantly for variations in parameters *k*_2_, *k*_4_, *k*_6_, *k*_8_, *k*_11_, *k*_20_, *k*_23_ and *k*_24_ [Supplementary Figure 1]. These parameters represent rates of: disassociation of RIG-MAVS complex(*k*_2_), disassociation of RIG-MAVS-Poly I:C complex (*k*_4_), association of RIG-MAVS-Poly I:C complex with TRAF-3 (*k*_6_), association of MAVS-RIG-Poly I:C-TRAF3 complex and TBK-1 (*k*_8_), dephosphorylation of TBK-1 (*k*_11_), translocation of phophorylated IRF-3 monomer from nucleus to cytoplasm (*k*_20_), and degradation of RIG and MAVS (represented by *k*_23_ and *k*_24_, respectively). A slight increase in the level of nuclear and active IRF-3 would be observed with an increase in the value of parameters representing rates of: binding of RIG and MAVS (*k*_1_), phosphorylation of TBK-1 (*k*_5_), binding of phosphorylated TBK-1 and RIG-MAVS-Poly I:C complex (*k*_9_) and translocation of phosphorylated IRF-3 from cytoplasm to the nucleus (*k*_17_). A decrease in the rate *k*_1_, *k*_5_ and *k*_9_ leads to a significant decrease in the nuclear concentration of active IRF-3 [Supplementary Figure 1]. A significant increase and decrease in the active nuclear phosphorylated IRF-3 levels is observed for variations in the rate of association of Poly I:C and RIG-MAVS complex (*k*_3_). A significant reduction in the IRF-3 levels is observed on increase in the value of following model parameters: rate of disassociation of MAVS-RIG-Poly I:CTRAF-3 complex (*k*_7_), rates of disassociation of MAVS-RIG-Poly I:C-TRAF-3-TBK 1 (phosphorylated) complex (*k*_10_) and (*k*_13_), and rate of export of phosphorylated IRF-3 dimer from nucleus to the cytoplasm (*k*_18_). A slight decrease in the level of IRF-3 would be observed with an increase in the value of the parameter denoting the rate of disassociation of RIG-I-MAVS-Poly I:C-TRAF-3 complex (*k*_12_). Neglebible change in IRF-3 levels would be observed for decrease in the values of *k*_12_).

The impact of changes in the values of a few model parameters on the levels of nuclear and active IRF-3 are shown in Figure 3. A slight increase in the level of nuclear and active IRF-3, would be observed for an increase in the rate of phosphorylation of IRF-3 in the cytoplasm (*k*_14_) (Figure 3A). However, a significant decrease in the level of IRF-3 would be observed for a decrease in the value of this parameter. No significant changes in the level of nuclear and active IRF-3 would be observed for variations in the rates of dimerization and monomerization of IRF-3 in the cytoplasm (*k*_15_ and *k*_16_, respectively) (Figure 3B and C).

An increase in the rate of import of cytoplasmic phosphorylated IRF-3 dimer (*k*_17_) would result in a faster and a 2-fold increase in the level of nuclear and active IRF-3. The active and nuclear levels of IRF-3 would drop sharply for decreases in the value of this parameter (Figure 3D).

An increase in the rate of monomerization of phosphorylated and nuclear IRF-3 (*k*_21_) would lead to a significant decrease (2- to 3- fold) in the level of nuclear and active IRF-3 (Figure 3E). A decrease in the value of the parameter (*k*_18_) would not show significant changes in the level of active and nuclear IRF-3 with respect to the level of that for the default value of *k*_18_ denoted by a black colored line)(Figure 3E). A slight increase, as well as decrease, in the active and nuclear IRF-3 concentration would be observed for an increase and decrease in the rate of dimerization of IRF-3 (phosphorylated) in the nucleus (*k*_22_), respectively (Figure 3F).

### 3.2 Expression Patterns of IFN*β* Co-regulated by NF*κ*B and IRF-3

To investigate the co-regulation of IFN*β* by IRF-3 and NF*κ*B, we have modeled the binding of them to the respective promoter regions using a framework proposed in Equation 2. Also, two Poly I:C input signals, shaped by the half-lives and initial concentration of Poly I:C in human serum from an earlier study [53] were considered (Figure 4 A). In addition, the active concentration of NF*κ*B and IRF-3 (Figure 4B and 4C, respectively) inside the nucleus were considered as inputs in Equation 2.

Our results suggest an immediate and sharp increase in the level of nuclear NF*κ* B in response to Poly I:C input signals (Figure 4B). Rapidly, the level of nuclear NF*κ*B reaches a maximum at *≈* 50 minutes and after that, it starts to decrease. The activation of nuclear NF*κ* B lasts for about 100 minutes, coincident with the disappearance of the Poly I:C signal ensuring no antiviral and inflammatory response once the stimulatory signal vanishes. In contrast, delayed activation of nuclear and active IRF-3 would be observed post-exponential attenuation of the Poly I:C signal (Figure 4C). The activation of IRF-3 would initially increase monotonically and reach a plateau. Due to that, a sustained antiviral and inflammatory response would be observed for a longer period. We noticed that the increase in the level of active IRF-3 in the nucleus starts exactly when the Poly I:C signal and the nuclear concentration of NF*κ*B vanishes.

We have modeled the individual promoter binding activity of NF*κ*B and IRF-3 using Equation 1 and the expression pattern of IFN*β*, i.e., the level of transcribed mRNA was calculated. Our results suggest an immediate and strong IFN*β* response when only NF*κ*B is bound and IRF-3 is not bound (Figure 5A), and a delayed sigmoid-shaped IFN*β* response when only IRF-3 is bound and NF*κ*B is not bound (Figure 5B). A comparatively more realistic situation occurs when both NF*κ*B and IRF-3 binds to their respective binding sites in the promoter region upstream of IFN*β*, which we have modeled through Equation 2. Equation 1 is a probabilistic modeling framework for the binding of NF*κ*B and IRF-3. The model results suggest that an early peak followed by a decline in the level of IFN*β* along with a sustained level of that would be observed for a longer time (Figure 5C)). The peak would decline after a certain time (*≈* 100 min) with a subsequent increase in the level of IFN*β* albeit slightly lower than the height of the initial peak but would remain sustained for a longer duration.

**Figure 5:**
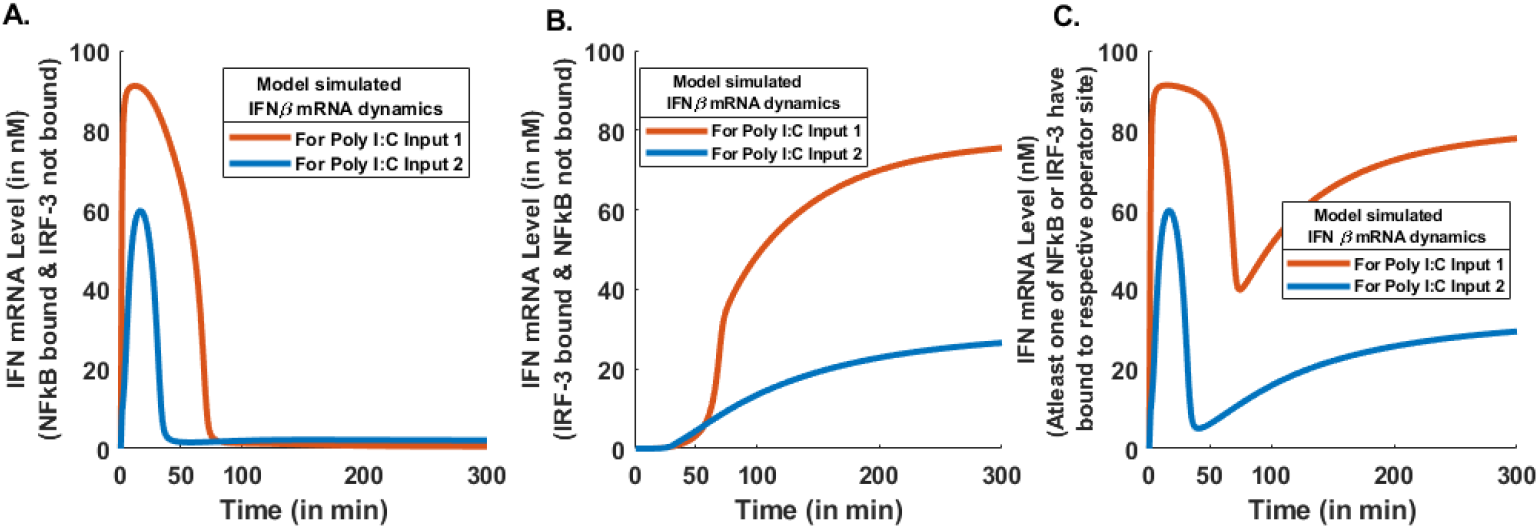
Expression pattern of IFNβ in response to input signal Poly I: C and mediated by either NFκB or IRF-3 as well as jointly. **(A.)** A pattern of IFNβ gene expression in response to Poly I:C Input 1 (orange curve) and Poly I:C Input 2 (blue curve) when NFκB binds to the promoter region and IRF-3 does not. It is modeled using a framework proposed in the Equation 2.**(B.)** Expression pattern of IFNβ gene in response to Poly I:C Input 1 (orange curve) and Poly I:C Input 2 (blue curve) when IRF-3 binds to its promoter region and NFκB does not.**(C.)** Change in the pattern of IFNβ expression in response to Poly I:C Input 1 (orange curve) and Poly I:C Input 2 (blue curve) when both NFκB and IRF-3 bind to their respective operator sites and show synergistic effects.

This expression pattern of IFN*β* is consistent with an observation for Newcastle Disease Virus (NDV) and Vesicular Stomatitis Virus (VSV) [34]. Wang et. al.(2010), in their experiments on mouse embryonic fibroblasts (MEFs) demonstrated that the replication of NDV and VSV were significantly higher in RelA^*−/−*^ compared to wild-type (WT) cells. The expression of IFN*β* in WT and RelA^*−/−*^ MEFs was similar at later times but substantially reduced in RelA^*−/−*^ MEFs at early time points (3-6 hours). This observation suggests a critical role of Rel A (a subunit of NF*κ*B) for initiating IFN*β* response. In addition, IRF-3 activation (determined by dimer formation) was not detected up to 6 hours after NDV infection by Western blot in WT and IRF-3 ^*−/−*^. At 12 hours and onwards, the active IRF-3 level reached a peak and the expression of IFN*β* was increased in WT but not in IRF-3 ^*−/−*^. Therefore, their results suggest that NF*κ*B is required for early antiviral response (i.e. increased level of IFN*β*) when IRF-3 activation is low, and IRF-3 alone is sufficient for elicitation of antiviral response during the late phase, consistent with our results.

We also calculated the IFN*β* response for two inputs of Poly I:C with different half-lives and for three binding scenarios (Figures 4A and 5). These results suggest that the IFN*β* response would not be exclusive to a particular viral analogue. Also, a stronger viral analogue (Poly I:C Input I) would lead to elevated levels of NF*κ*B, IRF-3 and IFN*β* inside the nucleus. A suppressed level of these species would be observed for Poly I:C input II (Figure 4,5).

### 3.3 Effect of Systematic Parameter Variations on the Expression Pattern of IFN*β*

We modeled the binding activity of both the transcription factors NF*κ*B and IRF-3 at the promoter sites of IFN*β* using a probabilistic framework (Equation 2) and estimated the level of expression of IFN*β* regulated by both transcription factors. In the absence of experimentally known values of the model parameters, we systematically varied model parameters by two and three orders of their magnitude to investigate the systems-level dynamic behavior (Figure 6). Poly I:C input I with a half-life of 20 minutes and initial concentration of 500 nM, was used as input for the present analysis of the model. Our results suggest that an increase of 2 or 3 orders of magnitude in the rate of phosphorylation of IRF-3 in the cytoplasm (*k*_14_) would lead to a slight increase in the expression of IFN*β* with respect to the level of that for the default value of *k*_14_ denoted by a black colored curve (Figure 6A). Post *≈* 100 minutes,the level of IFN*β* expression falls below the level of that for the default value of *k*_14_ for a 2 or 3 order of magnitude decrease in *k*_14_. A rapid decline after ≈ 100 minutes coincides with the rapid fall in Poly I:C concentration.

**Figure 6:**
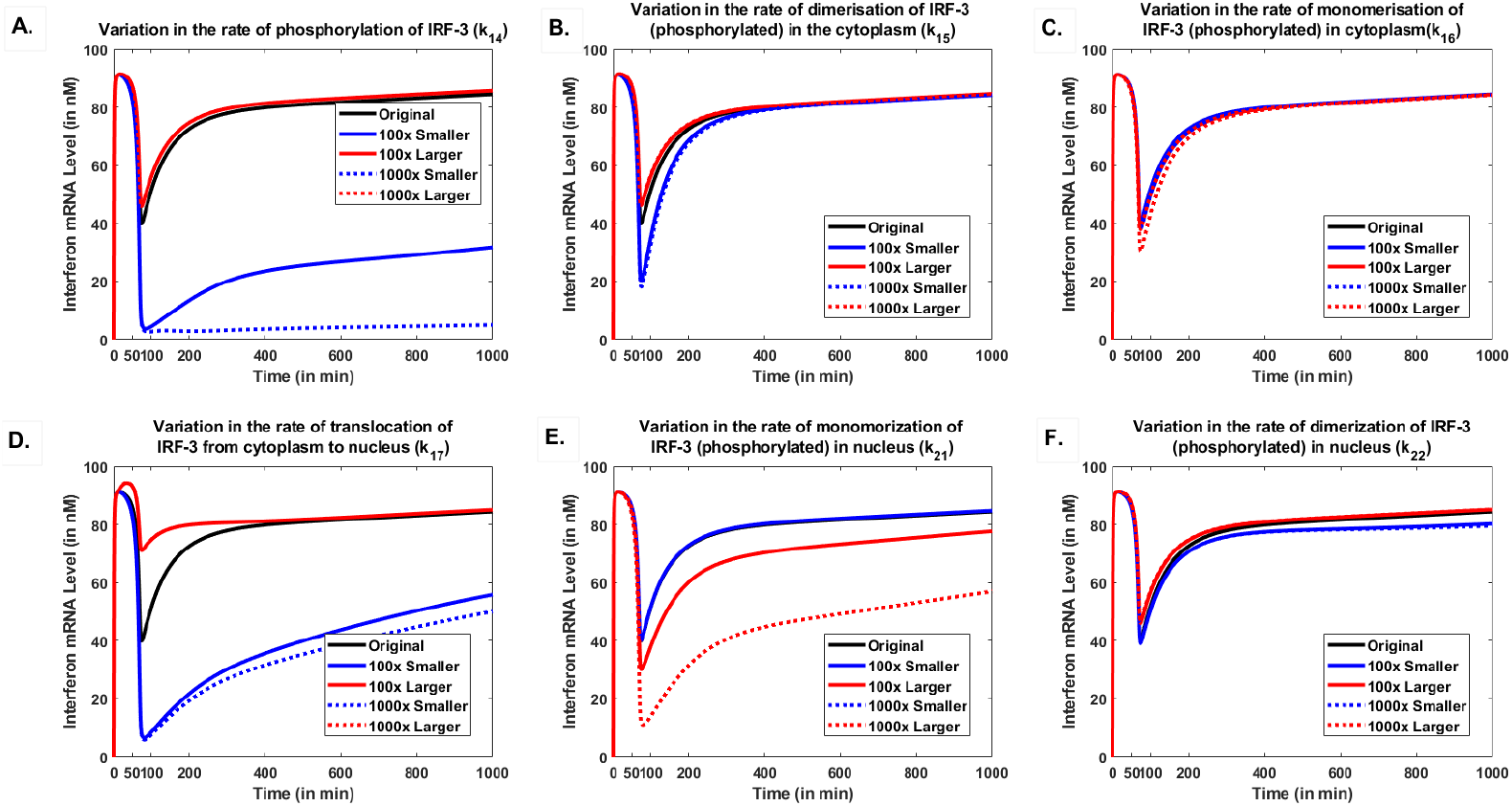
Effect of changes in model parameter’s values (k14 - k19, described in Table1,2) on the expression pattern of IFNβ over time. Blue colored lines represent the effect of decreases while red colored lines denote the effect of increase in the value of respective parameters by 2- or 3-orders of magnitude.

Furthermore, the results suggest that an increase in the rate of dimerization of IRF-3 (phosphorylated) in the cytoplasm (*k*_15_) of two or three orders of magnitude would lead to a slight increase in the expression of IFN*β* with respect to the expression of IFN*β* for the default value of *k*_15_ during 0 and 300 minutes (Figure 6B). A two-order or three order of magnitude decrease in the value of *k*_15_ would lead to a decrease in the level of IFN *β* for a short duration (100 to 300 minutes). After 300 minutes, the expression of IFN*β* would bounce back to the level of IFN *β* for the default of *k*_15_.

A negligible effect would be observed due to two or three orders of magnitude increase or decrease in the rate of monomerization of phosphorylated IRF-3 in the cytoplasm (*k*_16_) (Figure 6C).

An increase in the rate of translocation of the phosphorylated dimer of IRF-3 from the cytoplasm to the nucleus (*k*_17_), by two or three orders of magnitude, a sudden increase in the levels of IFN*β* would be observed between 0 to 50 minutes (Figure 6D). After *≈* 400 minutes, it approaches to a level of IFN*β* that would be observed for a default value of the parameter *k*_17_. For a two or three orders of magnitude reduction in *k*_17_ would result in a decrease in IFN*β* expression at 50 minutes and after that.

In addition, a decrease in the rate of monomerization of phosphorylated IRF-3 (*k*_21_) by two or three orders of magnitude would lead to a level of IFN*β* expression that would be observed for default parameter value of *k*_21_ (Figure 6E). During initial response (*≈* 50 minutes), there would be negligible effect of the change in the value of the parameter *k*_21_. However, an increase in *k*_21_ would lead to a drastic decrease in the level of IFN*β* expression after 100 minutes.

Also a two or three-orders of magnitude increase or decrease in the rate of dimerization of phosphorylated form of IRF-3 inside the nucleus (*k*_22_), would lead to a negligible change in the IFN*β* expression pattern (Figure 6F).

### 3.4 Role of the Binding Affinities of NF*κ*B and IRF-3 on the Expression Pattern of IFN*β*

Binding affinity is one of the measures to quantify the strength of interaction between tran-scription factors and their specific DNA binding sites at the promoter region upstream of a gene. Previous studies have proposed a mathematical framework to establish a relationship between binding affinity and its transcriptional output. The seminal study by Kim and O’Shea(2008) [44] demonstrates that strong binding affinity (or lower disassociation constant *k*_*d*_) leads to increased transcription of the target gene. However, a weak binding affinity (or a higher disassociation constant *k*_*d*_) leads to reduced transcription of the target gene. The general binding affinity of the transcription factor NF-*κ*B to its canonical sites *κ*B, reported as the equilibrium dissociation constant (*k*_*d*_), is 3–10 nM under physiological conditions (37 ^*°*^ C, 150 mM NaCl, and pH 7.5) replicating intracellular environments [54]. This range is not specific to the *κ*B site upstream of any particular type of IFN. However, the binding affinity of the transcription factor IRF-3 to the upstream of IFNs is not known.

In the present study, we investigate the effect of relative changes in the binding affinities of NF*κ*B and IRF-3 on the expression pattern of the inflammatory gene IFN*β*. We calculated the levels of transcribed mRNA levels for the following three distinct possibilities.

**Case I: k**_**NFkB**_ *>* **k**_**IRF***−***3**_. We kept the binding affinity for NF*κ*B based on the values reported in the literature [54]) (*k*_*NFkB*_ = 10 nM). The higher binding affinities (lower dissociation constants)for IRF-3 compared to that of the NF*κ*B were considered by decreasing *k*_*IRF* −3_ by a factor of 10, 100, and 1000, respectively.

**Case II:k**_**NFkB**_ *<* **k**_**IRF***−***3**_. The binding affinity of NF*κ*B was kept fixed through *k*_NF*κ*B_ = 10 nM and lower binding affinities (higher dissociation constants) for IRF-3 were considered by increasing *k*_*IRF* −3_ by a factor of 10, 100, and 1000, respectively.

**Case III : k**_**NFkB**_ = **k**_**IRF***−***3**_. The binding affinities of NF*κ*B and IRF-3 were kept equal. The same was varied within the observed range of the dissociation constant for NK*κ*B i.e. between 3 to 10 nM.

For Case I (*k*_*NFkB*_*>k*_*IRF* −3_), model results suggest that for a 10, 100, and 1000-fold increase in the binding affinity of IRF-3, the IFN*β* expression level would increase rapidly (Figure 7A). For all values of *k*_*IRF* −3_, we observed a maximum increase at *≈* 50 minutes. After that, it drops and settles at *≈* 90 nM. We noticed that the early response remains the same for all the values of *k*_*IRF* −3_, and the expression level of IFN*β* changes only after 100 minutes when the role of IRF-3 sets in. These results suggest that the higher binding affinity of IRF-3 would be responsible for stronger antiviral responses. For Case II (*k*_*NFkB*_ *< k*_*IRF* −3_), our results suggest that for a 10, 100, and 1000-fold decrease in the binding affinity of IRF-3, the IFN*β* expression level would increase rapidly initially followed by a rapid decline after 100 minutes (Figure 7B). Here, a 10-fold decrease in the binding affinity of IRF-3 would lead to a decrease in the transcribed IFN*β* mRNA level and settle at *≈* 20 nM. A 100 and 1000-fold decrease in the value of *k*_*IRF* −3_ would result in a further decrease in the final IFN*β* mRNA levels (*≈* 10 nM). Since the binding affinity of IRF-3 is weaker than that of NF*κ*B, it does not sustain a high level of transcribed IFN*β* mRNA after 100 minutes. In Case III when *k*_*NFkB*_ = *k*_*IRF* −3_, model results suggest the antiviral response would be greater than Case II but less than Case I (Figure 7C). A rapid increase in the expression level of IFN*β* would be observed followed by a slight dip around 100 minutes. The overall levels of transcribed IFN*β* mRNA would decrease with increasing values of *k*_*NFkB*_ and *k*_*IRF* −3_. In this case, when both the binding affinities of NF*κ*B and IRF-3 are equal, both NF*κ*B and IRF-3 transcription factors would be responsible for early antiviral response (IFN*β* 90 nM) and sustained (between 70 to *≈* 90 nM) for a longer period. These results collectively suggest that the binding affinity of IRF-3 would be either equal to or larger than that of NFkB (*k*_*NFkB*_ *≥k*_*IRF* −3_) for a strong and sustained antiviral response (Figure 7).

**Figure 7:**
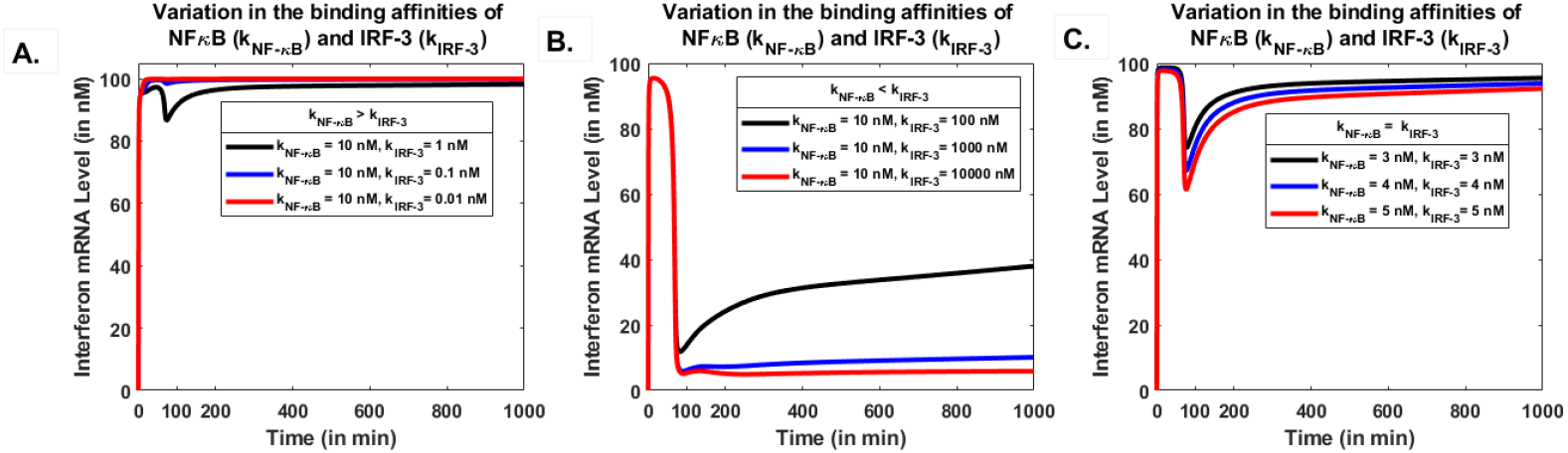
Role of the binding affinities of NF-κB (k_NF-κB_) and IRF-3 (k_IRF-3_) in the co-regulation of IFNβ. Blue colored lines represent the effect of decreases, while red colored lines denote increases in the respective parameters by 2 or 3 orders of magnitude. (A) IFNβ mRNA expression levels over time for k_NF-κB_ = 10 nM and varying k_IRF-3_: 1 nM (blue), 0.1 nM (orange), and 0.01 nM (yellow). (B) IFNβ levels for k_NF-κB_ = 10 nM and k_IRF-3_ of 10 nM (blue), 100 nM (orange), and 1000 nM (yellow). (C) IFNβ expression for equal binding affinities of k_NF-κB_ and k_IRF-3_: 3 nM (blue), 4 nM (orange), and 5 nM (yellow).

## 4 Discussion

Inflammation and tissue damage is a common phenomenon observed across diseases such as viral infection [55] and autoimmune disorders [56] as well as a part of a protective mechanism of the immune system [57]. It is widely accepted that inflammation induced by infections and tissue damage is an essential mechanism of innate immune response. Proper inflammatory responses provide broad-spectrum protection against infections and orchestrate long-term adaptive immunity towards specific pathogens. One of the mechanisms of inflammation is the expression of inflammatory genes such as type I IFNs, ISGs, IL-1*β*, IL-18, and IL-10 [58]. The activation of pattern recognition receptors and production of effector molecules are essential components of the innate immune system following viral infections [59]. Nuclear Factor *κ* B is one of the transcription factors that have a role in the expression of inflammatory genes in multiple diseases [15]. It has been established that the canonical NF*κ*B pathway has a role in eliciting the IFN response, which is one of the first lines of defense in the host’s immune system [60, 28, 61]. However, the type I IFN response is regulated by more than one signaling pathway for example by the transcription factor IRF-3 along with NF*κ*B post-viral infection. Viral input to the host’s cells is sensed by pattern recognition receptors (PRRs) such as TLRs (Toll-like receptors) and RLR (RIG like receptors) that activate the NF*κ*B and IRF-3 signaling pathways [62]. A study utilizing CHIP-seq and Genomewide association analysis (GWAS) has established that both the transcription factors NF*κ*B and IRF-3 cooperate to recruit factors that initiate the transcriptional machinery of IFN genes [63]. It has been computationally predicted that the promoter of IFN*β* contains NF-*κ*B binding sites and two IFN-stimulated responsive elements (ISREs) recognized by phosphorylated IRF3/7 [60]. Thus suggesting the role of both the transcription factors in co-regulating the expression of IFN*β*.

Previously, the induction of IFN*β*s by NF*κ*B and IRF-3 signaling pathways has been modeled along with their negative regulators, although using a stochastic approach [28]. There is no mathematical model available to compute the IFN*β* expression pattern in response to the viral analog, Poly I:C which is attempted in the present study.

Here, we have developed a mathematical model for the co-regulation of the IFN*β* gene by NF*κ*B and IRF-3 signaling pathways. The canonical signaling arm of NF*κ*B and IRF-3 was modeled using a mathematical model based on a deterministic ODE framework published elsewhere [27]. It is believed that the canonical NF*κ*B pathway plays a role in the elicitation of IFN*β* response [64, 65, 66], therefore, we gave null input to the NIK enzyme that activates the non-canonical NF*κ*B pathway. We modeled the IRF-3 signaling pathway by systematically identifying all the biochemical species and reactions involved, based on the existing literature [28]. A systematic parameter variation analysis by varying model parameters of the model of IRF-3 pathway was performed to explore the robustness of the model outputs and to identify critical parameters.

The DNA-NF*κ*B and DNA-IRF-3 binding are modeled here using a Hill-like equation to elicit the role of active NF*κ*B and IRF-3 in the regulation of IFN*β* expression. The level of the transcribed mRNA of IFN*β* was calculated using the concentration of respective transcription factors within the nucleus. In addition, we explored the expression pattern of IFN*β* for three scenarios: when (i) only NF*κ*B (ii) only IRF-3 and (iii) both NF*κ*B and IRF-3 bind the respective binding sites upstream to IFN*β*.

Our results suggest that a peak is formed in the response curve of the NF*κ*B lasting until 100 minutes (*≈* 1 to 2 hours) whereas significant active IRF-3 level appears only after *≈* 70 minutes. Therefore, the expression pattern of transcribed IFN*β* mRNA level would reach a maximum immediately presumably due to the binding of NF*κ*B and post 100 minutes due to the binding of IRF-3 to the binding sites in the upstream region of IFN*β*. Similar patterns of IRF-3 and NF*κ*B activation are observed in studies for NDV and VSV viral infections performed in MEFs [34].

In addition, these results suggest that the expression of IFN*β* is coregulated by the activation of NF*κ*B and IRF-3 at different time points in response to the viral analogue Poly I:C. Therefore, we propose a temporal separation in the activity of NF*κ*B and IRF-3 transcription factors. This is consistent with an earlier hypothesis of sequential transcriptional control proposed by Cheng et al. (2017) [35]. Through a transcriptomic and logic-gate based modeling approach, Cheng et al. have demonstrated that the expression of IFN *β* is regulated by temporal-separation in the activity of transcription factors, rather than their simultaneous co-binding. They have identified gene clusters regulated by either NF*κ*B or ISGF3 (an IRF-3 family member) or both, and proposed “sequential OR” and “sequential AND” logic gates based on the difference in transcriptional timing. We argue that a rapid and strong response due to NF*κ*B followed by a sustained response due to IRF-3 resembles a “sequential OR” gate mentioned by Cheng et al.

Further, the effect of the relative change in the binding affinities of the transcription factors NF*κ*B and IRF-3 on the transcribed level of IFN*β* are studied. Our results suggest that the binding affinity of IRF-3 would be equal to or larger than that of NF*κ*B (i.e. for dissociation constant, *k*_*NFkB*_ *≥ k*_*IRF* −3_) for a strong and sustained antiviral response.

One of the limitations of our model is that we have not considered the positive and negative feedback regulators of the biochemical species involved in the IRF-3 signaling pathway [42], or novel regulators of Type I IFN induction that are activated along with IRF-3 [44]. In addition, the present model does not consider the multiscale and multi-tissue dynamics which are considered in some of the stochastic mathematical models [45].

In future studies, the present model can be extended by considering the negative regulators of the IRF-3 signaling pathway. The construction of any mathematical model is constrained by the lack of experimental data, and the parameter estimation remains a fundamental challenge. The future models would addressed theses challenges to build a more pricise model .

## Declarations

## Acknowledgments

ST acknowledges the Prime Minister Research Fellowship awarded by the Ministry of Education, Government of India for financial assistance. RP acknowledges the Department of Science and Technology, India, for the DST-INSPIRE Faculty Award (DST/INSPIRE/04/2015/001939) and Banaras Hindu University for the Institute of Eminence seed grant. RP also acknowledges the University Grant Commission, India, for the start-up grant awarded to him.

## References

[1] I. Rahman and W. MacNee, “Role of transcription factors in inflammatory lung diseases,” Thorax, vol. 53, no. 7, pp. 601–612, 1998.

[2] E. Zaslavsky, U. Hershberg, J. Seto, A. M. Pham, S. Marquez, J. L. Duke, J. G. Wetmur, B. R. tenOever, S. C. Sealfon, and S. H. Kleinstein, “Antiviral response dictated by choreographed cascade of transcription factors,” Journal of Immunology, vol. 184, pp. 2908–2917, Mar 2010.

[3] P. Sengupta and S. Chattopadhyay, “Interferons in viral infections,” Viruses, vol. 16, p. 451, Mar 2024.

[4] L. Dalskov, H. H. Gad, and R. Hartmann, “Viral recognition and the antiviral interferon response,” EMBO Journal, vol. 42, no. 14, p. e112907, 2023. Epub 2023 Jun 27.

[5] F. McNab, K. Mayer-Barber, A. Sher, et al., “Type I interferons in infectious disease,” Nature Reviews Immunology, vol. 15, pp. 87–103, 2015. Published 23 January 2015, Issue Date: February 2015.

[6] A. I. Wells and C. B. Coyne, “Type III Interferons in antiviral defenses at barrier surfaces,” Trends in Immunology, vol. 39, pp. 848–858, Oct. 2018. Epub 2018 Sep 12.

[7] R. Hartmann, C. Weininger, O. V. Dontsova, and P. Kovarik, “Viral recognition and the antiviral interferon response,” The EMBO Journal, vol. 42, no. 15, p. e112907, 2023.

[8] K. Song and S. Li, “The role of ubiquitination in NF-κB signaling during virus infection,” Viruses, vol. 13, no. 2, p. 145, 2021.

[9] H. Yu, L. Lin, Z. Zhang, et al., “Targeting NF-κB pathway for the therapy of diseases: mechanism and clinical study,” Signal Transduction and Targeted Therapy, vol. 5, p. 209, 2020.

[10] K. Peters, H. Smith, G. Stark, and G. Sen, “IRF-3-dependent, NFκB-and JNK-independent activation of the 561 and IFN-β genes in response to double-stranded RNA,” Proceedings of the National Academy of Sciences of the United States of America, vol. 99, no. 9, pp. 6322–6327, 2002.

[11] B. Weaver, K. Kumar, and N. Reich, “Interferon regulatory factor 3 and CREB-binding protein/p300 are subunits of double-stranded RNA-activated transcription factor DRAF1,” Molecular and Cellular Biology, vol. 18, pp. 1359–1368, Mar 1998.

[12] M. Bao, N. Hofsink, and T. Plösch, “LPS versus Poly I:C model: comparison of long-term effects of bacterial and viral maternal immune activation on the offspring,” Am J Physiol Regul Integr Comp Physiol, vol. 322, no. 2, pp. R99–R111, 2022.

[13] D. Schweinoch, P. Bachmann, D. Clausznitzer, M. Binder, and L. Kaderali, “Mechanistic modeling explains the dsrna length-dependent activation of the RIG-I mediated immune response,” Journal of Theoretical Biology, vol. 500, p. 110336, 2020. Accessed: 2024-12-19.

[14] W. Zhu, J. Li, R. Zhang, Y. Cai, C. Wang, S. Qi, S. Chen, X. Liang, N. Qi, and F. Hou, “TRAF-3 IP-3 mediates the recruitment of TRAF-3 to MAVS for antiviral innate immunity,” The EMBO Journal, vol. 38, no. 18, p. e102075, 2019.

[15] T. Liu, L. Zhang, D. Joo, et al., “NFκB signaling in inflammation,” Sig Transduct Target Ther, vol. 2, no. 1, p. 17023, 2017.

[16] T. Jing, B. Zhao, P. Xu, X. Gao, L. Chi, H. Han, B. Sankaran, and P. Li, “The structural basis of IRF-3 activation upon phosphorylation,” J Immunol, vol. 205, no. 7, pp. 1886–1896, 2020.

[17] T. M. Petro, “IFN Regulatory Factor-3 in health and disease,” J. Immunol., vol. 205, no. 8, pp. 1981–1989, 2020.

[18] A. García-Sastre and C. A. Biron, “Type I Interferons and the virus-host relationship: a lesson in détente,” Science, vol. 312, pp. 879–882, May 2006.

[19] M. J. Clemens, “Translational control in virus-infected cells: models for cellular stress responses,” Seminars in Cell Developmental Biology, vol. 16, pp. 13–20, feb 2005.

[20] S. Martens and J. Howard, “The interferon-inducible GTPases,” Annual Review of Cell and Developmental Biology, vol. 22, no. 1, pp. 559–589, 2006.

[21] O. Haller, P. Staeheli, and G. Kochs, “Interferon-induced Mx proteins in antiviral host defense,” Biochimie, vol. 89, no. 6-7, pp. 812–818, 2007.

[22] Z. Talloczy, W. Jiang, H. W. Virgin IV, D. A. Leib, D. Scheuner, R. J. Kaufman, E.-L. Eskelinen, and B. Levine, “Regulation of starvation-and virus-induced autophagy by the eIF2α kinase signaling pathway,” Proceedings of the National Academy of Sciences, vol. 99, no. 1, pp. 190–195, 2002.

[23] Z. Tallóczy, H. Virgin IV, and B. Levine, “PKR-dependent xenophagic degradation of herpes simplex virus type 1,” Autophagy, vol. 2, no. 1, pp. 24–29, 2006.

[24] L. Espert, P. Codogno, and M. Biard-Piechaczyk, “Involvement of autophagy in viral infections: antiviral function and subversion by viruses,” Journal of Molecular Medicine, vol. 85, no. 8, pp. 811–823, 2007.

[25] G. Kochs, M. Trost, C. Janzen, and O. Haller, “MxA GTPase: oligomerization and GTP-dependent interaction with viral rnp target structures,” Methods, vol. 15, no. 3, pp. 255–263, 1998.

[26] F. Weber, O. Haller, and G. Kochs, “Mxa GTPase blocks reporter gene expression of reconstituted thogoto virus ribonucleoprotein complexes,” Journal of Virology, vol. 74, no. 1, pp. 560–563, 2000.

[27] T. Mukherjee, B. Chatterjee, A. Dhar, S. Bais, M. Chawla, M. Chawla, P. Roy, A. George, V. Bal, S. Rath, and S. Basak, “A TNF-p100 pathway subverts noncanonical NF-κB signaling in inflamed secondary lymphoid organs.,” The EMBO Journal, vol. 36, no. 23, pp. 3501–3516, 2017.

[28] M. Czerkies, Z. Korwek, W. Prus, M. Kochanczyk, J. Jaruszewicz-Blonska, K. Tudelska, S. Blonski, M. Kimmel, A. Brasier, and T. Lipniacki, “Cell fate in antiviral response arises in the crosstalk of IRF, NF-kB and JAK/STAT pathways,” Nature communications, vol. 9, no. 1, p. 493, 2018.

[29] R. Cheong, A. Hoffmann, and A. Levchenko, “Understanding NFκB signaling via mathematical modeling,” Mol Syst Biol, vol. 4, p. 192, 2008.

[30] J. Jaruszewicz-B lońska, I. Kosiuk, W. Prus, and T. Lipniacki, “A plausible identifiable model of the canonical NFκB signaling pathway,” PLoS ONE, vol. 18, no. 6, p. e0286416, 2023.

[31] S. Basak, M. Behar, and A. Hoffmann, “Lessons from mathematically modeling the NFκB pathway,” Immunological Reviews, vol. 246, pp. 221–238, Mar 2012.

[32] R. Williams, J. Timmis, and E. Qwarnstrom, “Computational models of the NFκB signalling pathway,” Computation, vol. 2, no. 4, pp. 131–158, 2014.

[33] T. Lipniacki, P. Paszek, A. R. Brasier, B. Luxon, and M. Kimmel, “Mathematical model of NFκB regulatory module,” Journal of Theoretical Biology, vol. 228, no. 2, pp. 195–215, 2004.

[34] J. Wang, S. H. Basagoudanavar, X. Wang, E. Hopewell, R. Albrecht, A. García-Sastre, and A. A. Beg, “NF-κB RelA subunit is crucial for early IFNβ expression and resistance to rna virus replication,” The Journal of Immunology, vol. 185, no. 3, pp. 1720–1729, 2010.

[35] Q. Cheng, A. Behzadi, N. Sen, Z. Ouyang, G. Ghosh, and A. Hoffmann, “Kinetic signaling pathways orchestrate sequential transcriptional logic in innate immunity,” Nature Immunology, vol. 18, no. 3, pp. 274–283, 2017.

[36] M. S. Hayden and S. Ghosh, “Crosstalk via the NF-κB signaling system,” Nature Reviews Immunology, vol. 6, no. 1, pp. 1–14, 2006.

[37] A. Hoffmann, A. Levchenko, M. L. Scott, and D. Baltimore, “Lessons from mathematically modeling the NF-κB pathway,” Cell, vol. 150, no. 4, pp. 707–716, 2012.

[38] J. D. Kearns, S. Basak, S. L. Werner, C. S. Huang, and A. Hoffmann, “Ib provides negative feedback to control NF-κB oscillations, signaling dynamics, and inflammatory gene expression,” The Journal of Cell Biology, vol. 173, no. 5, pp. 659–664, 2006.

[39] T. Sato, T. Shimozato, H. Nakano, K. Takeda, and S. Akira, “Coordination between NF-κB family members p50 and p52 is essential for mediating LTβR signals in the development and organization of secondary lymphoid tissues,” Journal of Immunology, vol. 177, no. 8, pp. 5332– 5340, 2006.

[40] M. Chawla, P. Roy, and S. Basak, “Role of the NF-κB system in context-specific tuning of the inflammatory gene response,” Current Opinion in Immunology, vol. 68, pp. 21–27, 2021.

[41] I. The MathWorks, MATLAB, 2023. R2023a.

[42] W. Zhang and X. Zou, “Systematic analysis of the mechanisms of virus-triggered Type I IFN signaling pathways through mathematical modeling,” IEEE/ACM Transactions on Computational Biology and Bioinformatics, vol. 10, no. 3, pp. 771–779, 2013.

[43] C. Cai and X. Yu, “A mathematic model to reveal delicate cross-regulation between MAVS/STING, inflammasome and MyD88-dependent type I interferon signalling,” Journal of Cellular and Molecular Medicine, vol. 24, no. 19, pp. 11535–11545, 2020.

[44] H. Kim and B. Seed, “The transcription factor MafB antagonizes antiviral responses by blocking recruitment of coactivators to the transcription factor IRF3,” Nature Immunology, vol. 11, no. 8, pp. 743–750, 2010.

[45] D. R. Ourthiague, H. Birnbaum, N. Ortenlöf, J. D. Vargas, R. Wollman, and A. Hoffmann, “Limited specificity of IRF3 and ISGF3 in the transcriptional innate-immune response to double-stranded rna,” Journal of Leukocyte Biology, vol. 98, no. 1, pp. 119–128, 2015.

[46] A. Günel, “Modelling the interactions between TLR4 and IFNβ pathways,” Journal of Theoretical Biology, vol. 307, pp. 137–148, 2012.

[47] X. Zou, X. Xiang, Y. Chen, and Z. Pan, “Mathematical modeling of signaling pathways leading to Type I IFN Gene Expression,” 2009.

[48] Z. Korwek et al., “Nonself rna rewires IFN-β signaling: A mathematical model of the innate immune response,” Science Signaling, vol. 16, no. 797, p. eabq1173, 2023.

[49] X. Zou, X. Xiang, Y. Chen, T. Peng, X. Luo, and Z. Pan, “Understanding inhibition of viral proteins on Type I IFN signaling pathways with modeling and optimization,” Journal of Theoretical Biology, vol. 265, no. 4, pp. 691–703, 2010.

[50] L. Bintu, N. Buchler, H. Garcia, U. Gerland, T. Hwa, J. Kondev, and R. Phillips, “Transcriptional regulation by the numbers: models,” Current Opinion in Genetics & Development, vol. 15, no. 2, pp. 116–124, 2005.

[51] U. Alon, An Introduction to Systems Biology: Design Principles of Biological Circuits. London: Chapman and Hall/CRC, 1 ed., 2006.

[52] N. E. Buchler, U. Gerland, and T. Hwa, “On schemes of combinatorial trancription logic,” Proceedings of the National Academy of Sciences, vol. 100, no. 9, pp. 5136–5141, 2003.

[53] E. de Clercq, “Degradation of Poly(inosinic acid).poly(cytidylic acid)[(I)n.(Cn)] by Human Plasma,” European Journal of Biochemistry, vol. 93, no. 1, pp. 165–172, 1979.

[54] S. Bergqvist, V. Alverdi, B. Mengel, A. Hoffmann, G. Ghosh, and E. Komives, “Kinetic enhancement of NF-κB·DNA dissociation by IκBα,” Proceedings of the National Academy of Sciences of the United States of America, vol. 106, no. 46, pp. 19328–19333, 2009.

[55] A. Faist, J. Janowski, S. Kumar, S. Hinse, D.M. Çalişkan, J. Lange, S. Ludwig, and L. Brunotte, “Virus infection and systemic inflammation: Lessons learnt from COVID-19 and beyond,” Cells, vol. 11, no. 14, p. 2198, 2022. Accessed: 2025-01-20.

[56] Y. Xiang, M. Zhang, D. Jiang, Q. Su, and J. Shi, “The role of inflammation in autoimmune disease: a therapeutic target,” Frontiers in Immunology, vol. 14, p. 1267091, Oct 2023.

[57] L. Chen, H. Deng, H. Cui, J. Fang, Z. Zuo, J. Deng, Y. Li, X. Wang, and L. Zhao, “Inflammatory responses and inflammation-associated diseases in organs,” Oncotarget, vol. 9, pp. 7204–7218, Dec 2017.

[58] M. H. Christensen and S. R. Paludan, “Viral evasion of DNA-stimulated innate immune responses,” Cellular & Molecular Immunology, vol. 14, no. 1, pp. 4–13, 2017.

[59] T. S. Xiao, “Innate immunity and inflammation,” Cellular & Molecular Immunology, vol. 14, no. 1, pp. 1–3, 2017.

[60] M. Iwanaszko and M. Kimmel, “NFκB and IRF pathways: cross-regulation on target genes promoter level,” BMC Genomics, vol. 16, pp. 1–8, 2015.

[61] L. C. Van Eyndhoven, A. Singh, and J. Tel, “Decoding the dynamics of multilayered stochastic antiviral IFN-I responses,” Trends in Immunology, vol. 42, no. 9, pp. 824–839, 2021.

[62] R. Bertolusso, B. Tian, Y. Zhao, L. Vergara, A. Sabree, M. Iwanaszko, T. Lipniacki, A. R. Brasier, and M. Kimmel, “Dynamic cross talk model of the epithelial innate immune response to double-stranded rna stimulation: Coordinated dynamics emerging from cell-level noise,” PLoS ONE, vol. 9, no. 4, p. e93396, 2014.

[63] J. E. Freaney, R. Kim, R. Mandhana, and C. M. Horvath, “Extensive cooperation of immune master regulators IRF3 and NFκB in RNA Pol II recruitment and pause release in human innate antiviral transcription,” Cell Reports, vol. 4, no. 5, pp. 959–973, 2013.

[64] S. Namineni, T. O’Connor, S. Faure-Dupuy, P. Johansen, T. Riedl, K. Liu, H. Xu, et al., “A dual role for hepatocyte-intrinsic canonical NF-κB signaling in virus control,” Journal of Hepatology, vol. 72, no. 5, pp. 960–975, 2020.

[65] M. J. Lenardo, C. M. Fan, T. Maniatis, and D. Baltimore, “The involvement of NF-κB in IFNβ gene regulation reveals its role as widely inducible mediator of signal transduction,” Cell, vol. 57, pp. 287–294, 1989.

[66] K. V. Visvanathan and S. Goodbourn, “Double-stranded RNA activates binding of NF-κB to an inducible element in the human beta-interferon promoter,” EMBO Journal, vol. 8, pp. 1129– 1138, 1989.

