## Supplementary File for "Modeling the Co-Activation of Inflammatory Genes Mediated by NFκB and IRF-3 During Viral Infections"

### Supplementary Information

May 13, 2025

#### Supplementary Tables & Figures

Table 1: **List of biochemical species (Model State Variables) in the IRF-3 Model**

| Serial No. | Biochemical Species | Model State Variable | Description |
|---|---|---|---|
| 0 | Poly I:C | POLYIC | Viral analogue |
| 1 | RIG | RIG | RIG-I protein |
| 2 | MAVS | MAVS | MAVS protein |
| 3 | MAVS:RIG | C1 | Complex |
| 4 | MAVS:RIG:PolyIC | C2 | Complex |
| 5 | TRAF3 | TRAF | TRAF3 protein |
| 6 | MAVS:RIG:PolyIC:TRAF3 | C3 | Complex |
| 7 | TBK1 | TBK | Protein |
| 8 | Phosphorylated TBK1 | TBK(p) | Protein |
| 9 | MAVS:RIG:PolyIC:TRAF3:TBK1(p) | C4 | Complex |
| 10 | $IRF3_{cytoplasm}$ | $IRF3_{c}$ | Protein |
| 11 | Phosphorylated $IRF3_{cytoplasm}$ | $IRF3_{p,c}$ | Protein |
| 12 | Phosphorylated $IRF3_{cytoplasm,dimer}$ | $IRF3_{p,c,d}$ | Protein |
| 13 | Phosphorylated $IRF3_{nuclear}$ | $IRF3_{p,n}$ | Protein |
| 14 | Phosphorylated $IRF3_{nuclear,dimer}$ | $IRF3_{p,n,d}$ | Protein |

Table 2: **List of initial concentrations in the IRF-3 Model**

| Serial No. | Biochemical Species | Concentration (nM) | Reference |
|---|---|---|---|
| 1 | RIG | 124.58 | Calculated from [1] |
| 2 | MAVS | 62.29 | Calculated from [1] |
| 3 | TBK | 124.58 | Calculated from [1] |
| 4 | TRAF | 62.29 | Calculated from [1] |
| 5 | IRFc | 124.58 | Calculated from [1] |

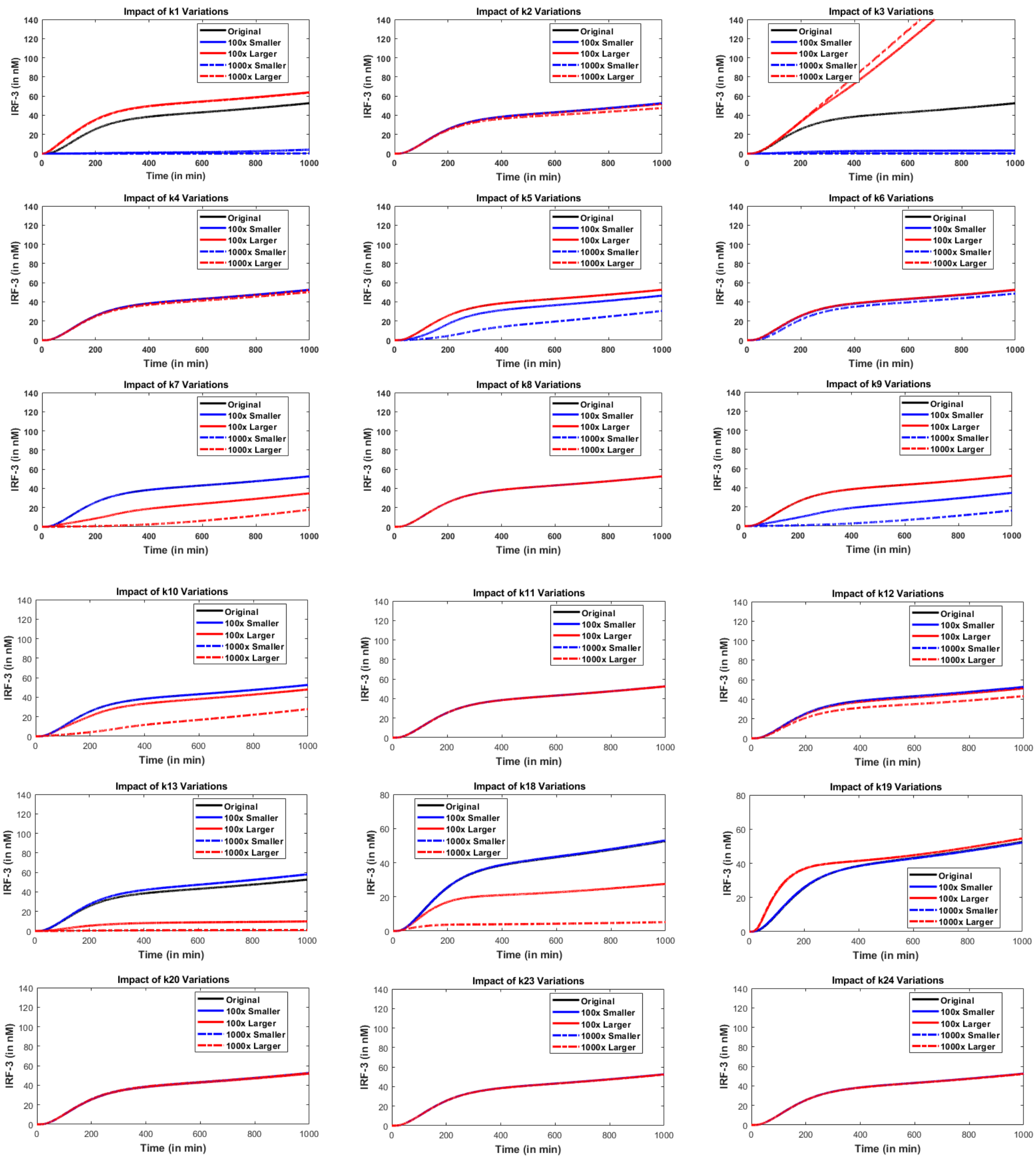


Figure 1: *Systematic parameter analysis showing the effect of varying model parameters* $k_1$ *to* $k_{13}$ *and* $k_{20}$ *to* $k_{24}$ *(as described in the main text, Table 1 and Table 2) on the expression of active IRF-3 levels in the nucleus.*

### System of ODEs Governing IRF-3 Signaling

The following set of ordinary differential equations describes the temporal behavior of molecular species involved in the IRF-3 signaling pathway in the IRF-3 Module. The model parameters ($k_1 - k_{29}$) of the ODE system are described in main text (Table 1 and Table 2).

#### ODE System

$$\frac{dy_1}{dt} = -k_1 y_1 y_2 + k_2 y_3 - k_{23} y_1 + k_{31} \tag{S1}$$

$$\frac{dy_2}{dt} = -k_1 y_1 y_2 + k_2 y_3 - k_{24} y_2 + k_{32} \tag{S2}$$

$$\frac{dy_3}{dt} = k_1 y_1 y_2 - k_2 y_3 - k_3 y_3 \cdot \text{POLYIC} + k_4 y_4 + k_{12} y_6 + k_{13} y_9 \tag{S3}$$

$$\frac{dy_4}{dt} = k_3 y_3 \cdot \text{POLYIC} - k_4 y_4 - k_6 y_4 y_5 + k_7 y_6 \tag{S4}$$

$$\frac{dy_5}{dt} = -k_6 y_4 y_5 + k_7 y_6 + k_{12} y_6 + k_{13} y_9 - k_{26} y_5 + k_{34} \tag{S5}$$

$$\frac{dy_6}{dt} = k_6 y_4 y_5 - k_7 y_6 - k_8 y_6 y_7 - k_9 y_8 y_6 + k_{10} y_9 - k_{12} y_6 \tag{S6}$$

$$\frac{dy_7}{dt} = -k_5 y_7 - k_8 y_7 y_6 + k_{11} y_8 - k_{25} y_7 + k_{33} \tag{S7}$$

$$\frac{dy_8}{dt} = k_5 y_7 - k_9 y_8 y_6 + k_{10} y_9 - k_{11} y_8 + k_{13} y_9 \tag{S8}$$

$$\frac{dy_9}{dt} = k_8 y_7 y_6 + k_9 y_8 y_6 - k_{10} y_9 - k_{13} y_9 - k_{14} y_{10} y_9 \tag{S9}$$

$$\frac{dy_{10}}{dt} = -k_{14} y_{10} y_9 - k_{27} y_{10} + k_{35} \tag{S10}$$

$$\frac{dy_{11}}{dt} = k_{14} y_{10} y_9 - 2k_{15} y_{11}^2 + 2k_{16} y_{12} - k_{19} y_{11} + k_{20} y_{13} \tag{S11}$$

$$\frac{dy_{12}}{dt} = k_{15} y_{11}^2 - k_{16} y_{12} - k_{17} y_{12} + k_{18} y_{14} - k_{29} y_{12} \tag{S12}$$

$$\frac{dy_{13}}{dt} = k_{19} y_{11} - k_{20} y_{13} + 2k_{21} y_{14} - 2k_{22} y_{13}^2 - k_{28} y_{13} \tag{S13}$$

$$\frac{dy_{14}}{dt} = k_{17} y_{12} - k_{18} y_{14} + k_{22} y_{13}^2 - k_{21} y_{14} - k_{30} y_{14} \tag{S14}$$

#### Biological Interpretation of Model State Variables

- $y_1$: RIG-I (pattern recognition receptor)
- $y_2$: MAVS (mitochondrial antiviral signaling protein)
- $y_3$: MAVS-RIG complex
- $y_4$: MAVS-RIG-PolyIC complex
- $y_5$: TRAF3 adapter
- $y_6$: MAVS-RIG-PolyIC-TRAF3 complex
- $y_7$: TBK1
- $y_8$: Phosphorylated TBK1
- $y_9$: MAVS-RIG-PolyIC-TRAF3-Phosphorylated TBK1 complex

- $y_{10}$: Cytoplasmic IRF-3 (inactive)
- $y_{11}$: Phosphorylated cytoplasmic IRF-3 monomer
- $y_{12}$: Phosphorylated cytoplasmic IRF-3 dimer
- $y_{13}$: Phosphorylated nuclear IRF-3 monomer
- $y_{14}$: Phosphorylated nuclear IRF-3 dimer (active)

### References

[1] M. Czerkies, Z. Korwek, W. Prus, M. Kochanczyk, J. Jaruszewicz-Blonska, K. Tudelska, S. Blonski, M. Kimmel, A.R. Brasier, and T. Lipniacki. Cell fate in antiviral response arises in the crosstalk of IRF, NF-kB and JAK/STAT pathways. *Nature communications*, 9(1):493, 2018.
